# Gammaretroviruses, novel viruses and pathogenic bacteria in Australian bats with neurological signs, pneumonia and skin lesions

**DOI:** 10.1101/2022.10.20.513122

**Authors:** Kate Van Brussel, Jackie E. Mahar, Jane Hall, Hannah Bender, Ayda Susana Ortiz-Baez, Wei-Shan Chang, Edward C. Holmes, Karrie Rose

## Abstract

More than 70 bat species are found in mainland Australia, including five species of megabat from a single genus (family Pteropodidae) and more than 65 species representing six families of microbats. The conservation status of these animals varies from least concern to endangered. Research directed at evaluating the impact of microorganisms on bat health has been generally restricted to surveillance for specific pathogens. While most of the current bat virome studies focus on sampling apparently healthy individuals, little is known about the infectome of diseased bats. We performed traditional diagnostic techniques and metatranscriptomic sequencing on tissue samples from 43 individual bats, comprising three flying fox and two microbat species experiencing a range of disease syndromes, including mass mortality, neurological signs, pneumonia and skin lesions. We identified reads from four pathogenic bacteria and two pathogenic fungi, including *Pseudomonas aeruginosa* in lung samples from flying foxes with peracute pneumonia, and with dermatitis. Of note, we identified the recently discovered Hervey pteropid gammaretrovirus, with evidence of replication consistent with an exogenous virus, in a bat with lymphoid leukemia. In addition, one novel picornavirus, at least three novel astroviruses and bat pegiviruses were identified. We suggest that the most likely cause of peracute lung disease was *Pseudomonas aeruginosa*, while we suspect Hervey pteropid gammaretrovirus was associated with lymphoid leukemia. It is possible that any of the novel astroviruses could have contributed to the presentation of skin lesions in individual microbats. This study highlights the importance of studying the role of microorganisms in bat health and conservation.

**IMPORTANCE:** Bats have been implicated as reservoir hosts for zoonotic disease of concern, however, the burden of microorganism including viruses on bat health and disease is understudied. Here we incorporated veterinary diagnostics and RNA sequencing to identify the presence of microbes and viruses with possible pathogenic status in Australian bats with varying disease presentations. These techniques were able to effectively identify and describe several pathogenic species of bacteria and fungi in addition to known and novel viruses. This study emphasises the importance of screening pathogens in cases of bat mortality for the conservation of this diverse order.

## INTRODUCTION

The mammalian order Chiroptera comprises over 1000 species of bat with a near global distribution. In recent years bats have gained attention for their ability to carry a large number of viruses, some of which have jumped hosts to emerge in new species (1). As the sampling of bats has increased dramatically over the last decade so the known bat virosphere has similarly expanded, including the discovery of numerous novel viruses in addition to new variants of existing zoonotic pathogens (2-5). Of particular importance is understanding the factors that enable bats to carry such a high diversity and abundance of viruses, likely reflecting unique immunological components in conjunction with such factors as large population densities and high body temperature during flight (6, 7). In turn, such research has led to a common belief that bats are able to tolerate a multitude of seemingly commensal viruses and do not experience large-scale outbreaks of infectious disease. Bats, however, are clearly susceptible to microbial infections with, for example, *Pseudogymnoascus destructans*, a fungus that causes white-nose syndrome, having a devastating effect on bats in North America (8, 9). In addition, infection with lyssaviruses can result in neurological and behavioural changes in bats (10).

Over 70 species of bat from the families Emballonuridae, Hipposideridae, Pteropodidae, Megadermatidae, Miniopteridae, Molossidae, Rhinolophidae, Rhinonycteridae and Vespertilionidae inhabit mainland Australia (11). As of 2021, the International Union for Conservation of Nature (IUCN) Red List of Threatened Species lists nine of these as vulnerable, six as near threatened, one as endangered (the spectacled flying fox, *Pteropus conspicillatus*) and two as extinct (11). Together with the endangered spectacled flying fox, three other *Pteropus* species inhabit Australia: the grey-headed flying fox (*P. poliocephalus*), listed as vulnerable, while the little red flying fox (*P. scapulatus*) and black flying fox (*P. alecto*) are listed as of least concern. The habitat range of these flying fox species includes the north and east of Australia, with the grey-headed flying fox inhabiting as far west as Adelaide, South Australia (12). Australia is also home to several insectivorous microbat species, which, together with flying foxes (frugivores and nectivores), play an essential role in maintaining Australia’s ecosystems by distributing seeds, pollinating plants and controlling insect numbers (13-15). Consequently, any major decline in bat numbers in Australia could have a negative impact on ecosystem health (15).

While mass mortalities of adult and young flying foxes have been associated with periods of extreme heat (16, 17), additional disease syndromes and mass mortality events have recently emerged in Australian chiropterans. Episodic mass pup abandonment has been associated with extreme heat, but also dehydration, nutritional stress, and dam death or desertion (16). Herein, we describe the emergence of several novel disease syndromes in flying foxes, including a distinctive pattern of acute to peracute vascular and inflammatory lung lesions in grey-headed flying foxes and a black flying fox following exposure to stressors such as extreme heat, mass pup abandonment, or traumatic injury. Additionally, an emergent syndrome of neurological disease in flying foxes is characterised by flaccid paralysis, severe central depression, tongue protrusion, and voice changes. Affected animals test negative for lyssavirus and often present thin, after periods of heavy rain. A dermatopathy in grey-headed flying foxes in extended rehabilitation care is characterised by depigmentation, ulceration and moist dermatitis of the wing webs. Individual cases in our study included wild flying-foxes with multisystemic lymphoma, and nodular wing web lesions associated with mite infestation.

To help identify the aetiological agents behind the presentation of severe disease in several bat species in Australia, we used traditional veterinary diagnostic techniques and a metatranscriptomic (i.e., total RNA sequencing) approach to characterise the histological change and viral and microbial diversity in tissues taken from bats displaying varying signs of disease, including neurological signs, peracute death and skin lesions (Table S1). The species of bat included in this study comprise grey-headed, black and little red flying foxes, as well as two species of microbats – eastern bent-wing bat (*Miniopterus orianae oceanensis)* and large footed myotis (*Myotis Macropus)* (Table S1).

## RESULTS

### Peracute to actue pneumonia in bats

Wildlife rehabilitators in the Sydney basin noted the rapid decline and demise of flying foxes following mild to moderate trauma. Affected animals refused food then became progressively weak, moribund and died within 12-48 hours. Post-mortem examination of these animals revealed a uniform pattern of voluminous lungs with multifocal petechial haemorrhages (Fig. 1a). Impression smears of affected lung tissue often contained fine bacilli, individually, forming palisades, or clustered within the cytoplasm of macrophages (Fig. 1a inset). Similar gross changes were noted in animals evaluated following extreme heat related mass mortalities or mass pup abandonment. On histologic examination of affected animal tissues, the lung lesions consisted of perivascular haemorrhage with interstitial oedema, fibrin deposition and necrosis (Fig. 1b). Lesions were often devoid of inflammatory cell infiltration (peracute), while others contained variable numbers of neutrophils and histiocytes (acute). *Pseudomonas aeruginosa* was isolated in microbial culture from nine of fifteen grey-headed flying fox lungs, and from a single black flying fox lung. Isolates were susceptible to a wide range of antimicrobial agents. Correlation between the identification of fine bacilli in lung impression smears with the isolation of *P. aeruginosa* was high. Although *P. aeruginosa* was the only isolate in six bat lung samples, *Escherichia coli, Klebsiella oxytoca, Lactococcus lactis, Enterobacter asburiae*, and *Streptococcus* species were also isolated in some lung tissues. *Salmonella enterica* serovar Wangata was also isolated in the lung and intestine of a young female grey-headed flying fox that died immediately after being rescued from a pup abandonment event. This animal had neutrophilic interstitial pneumonia, but also necrotising hepatitis, histiocytic colitis and evidence of septicaemia.

**FIG 1.**
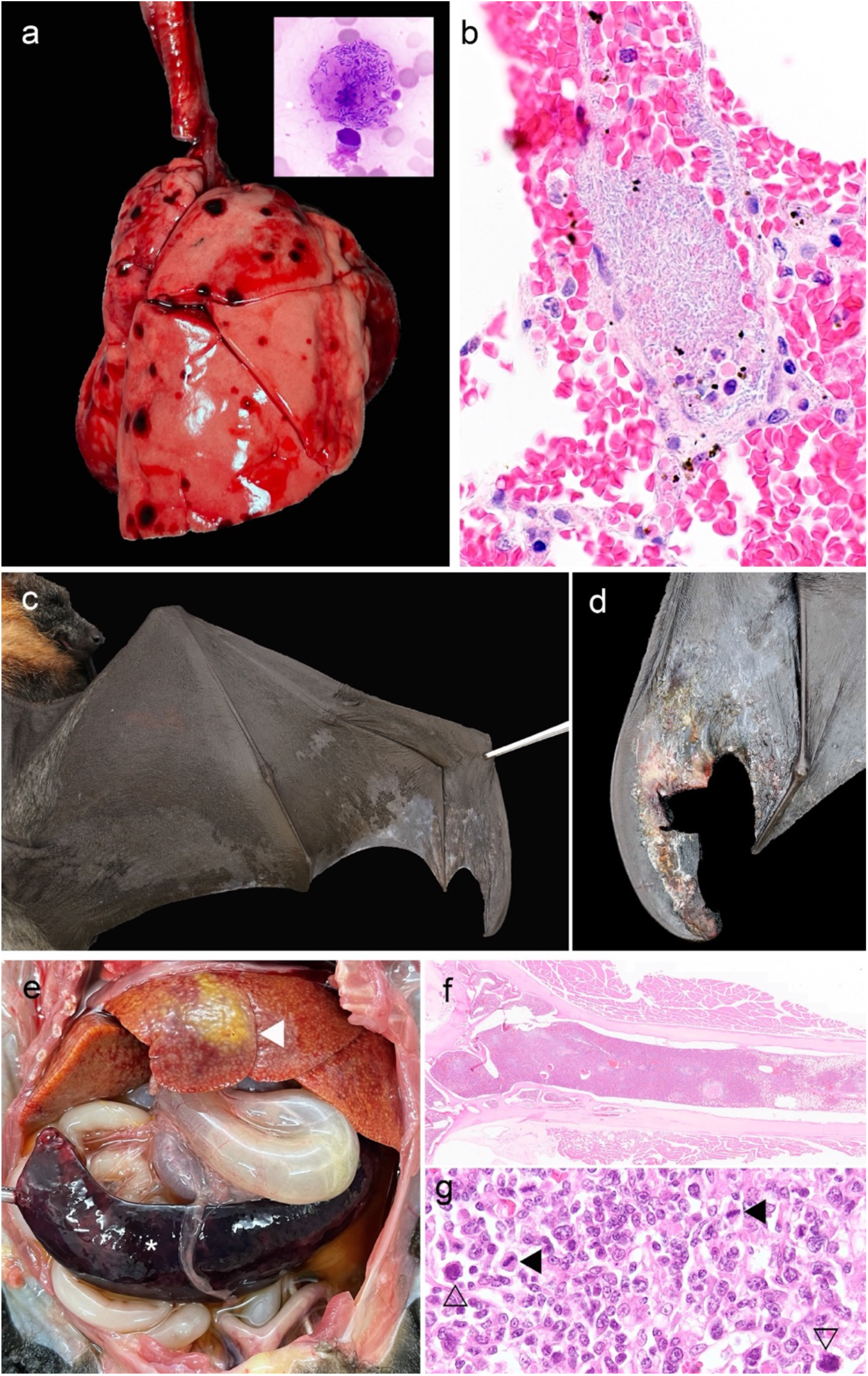
(a) Voluminous lungs with multifocal haemorrhages in a grey-headed flying fox. Inset: a pulmonary macrophage contains intracellular fine bacilli (lung impression smear, Quick Dip™ 1000x, case 14053.1). (b) Photomicrograph illustrating acute pulmonary perivascular haemorrhage associated with palisades of fine bacilli transmigrating an interstitial blood vessel wall. (c and d) Depigmentation and ulceration (d - with square excisional biopsy defect) extensively across the wing webs of grey-headed flying foxes (cases 14130.1 and 3). (e) Abdominal cavity of a grey-headed flying fox with abundant peritoneal fluid, a large liver with miliary, coalescing, raised, pale subcapsular foci (euthanasia artefact - white arrowhead) and marked splenomegaly (*). (f and g) Leukemia in a grey-headed flying fox. (f) The bone marrow is replaced with densely packed neoplastic cells (HE 20x). (g) Lymph node architecture is effaced by a confluence of medium sized round cells, some of which have bizarre nuclei (open arrowheads) or mitotic figures (black arrowheads) (case 14065.1, HE 1000x).

### Flying fox paralysis syndrome

An emergent, episodic syndrome of flaccid paralysis and central nervous system depression is characterised by bats that are recumbent, with protruding tongues, and unusual vocalisations. Affected bats can sometimes grasp on to wire or a branch: however, this is a passive action for the chiropteran foot, which can occur in the face of paralysis. Bats with this syndrome tend to present in clusters, or in large numbers (>150). Males are over-represented, and their body condition is generally poor, with weight 15-25% less than that expected based on the forearm measurement. Events tend to occur after periods of heavy rain. No significant histological lesions have been detected in affected animals, except for a single grey-headed flying fox that had lung lesions, as described above.

### Ulcerative and depigmenting dermatopathy in bats

Ecologists and wildlife rehabilitators often report patches of depigmentation of the flying fox wing web in free-ranging animals. Flying foxes in rehabilitation care have been observed to develop extensive depigmentation of the wing web with gross and histological evidence of hyperkeratosis, dermatitis and ulceration at the tips of the wing webs. Four affected young flying foxes in wildlife rehabilitation care were euthanised and examined (Fig. 1c and d). *Pseudomonas aeruginosa* was isolated from swabs collected from the active lesions of each animal. Although *P. aeruginosa* was a predominant isolate from each animal, moderate growth of *Serratia marcescens* was also grown in skin swabs from three animals. Variable growth of *Pseudomonas protegens, Enterococcus faecalis*, alpha haemolytic *Streptococcus*, and *Fusarium* species were also detected.

### Isolated bat cases

This study also included animals with no distinctive disease pattern. A single subadult male grey-headed flying fox with mite associated wing web lesions was euthanised after antiparasitic treatment resulted in central nervous system depression (bat no. 11501.1). A single adult female grey-headed flying fox was euthanised due to recumbency and marked abdominal distension (bat no. 14065.1). Post-mortem examination revealed marked peritoneal effusion with clear, straw-coloured fluid, and severe hepatosplenomegaly (Fig. 1e). The hepatic tissue contained a prominent zonal pallor, which appeared raised along the capsular surface. On histologic examination, the spleen, lymph nodes, bone marrow, periportal hepatic parenchyma, and portions of the kidney and adrenal gland were effaced by sheets of neoplastic cells (Fig. 1f). An impression smear of the spleen, and histological examination of affected tissues revealed a confluent array of monomorphic medium to large lymphocytes often exhibiting karyomegaly, reniform or bizarre nuclei, and two to three mitotic figures per 1000x field (1g), characteristic of lymphoid leukemia.

### Overview of metatranscriptomic data

In total, 32 tissue libraries were prepared for total RNA sequencing, comprising 10 from lung tissue, 10 from brain tissue, 10 from liver tissue and 2 from skin samples (Table S1). Overall, 38 grey-headed flying foxes, one black flying fox, two little red flying foxes, one eastern bent-wing bat and one large footed myotis from New South Wales (NSW) and the Australian Capital Territory (ACT) were included in this study (Table S1). Each sequencing library produced 87,000-253,000 reads and 112,000-696,000 contigs. Overall, we detected sequencing reads from 60 bacterial and 58 fungal families (Fig. S1). Additionally, we identified virus sequences belonging to 15 families, including novel viruses from the *Astroviridae* and *Picornaviridae* and known viruses from the *Flaviviridae* (pegivirus) and *Retroviridae* (gammaretrovirus).

### Overview of the bacteria and fungi present in bats

Although the bats sampled here had a diverse microbiome (Fig. S1), we detected high levels of read abundance for five potentially pathogenic bacteria and fungi in nine bat libraries: *Enterococcus faecalis, Pseudomonas aerugosina, Salmonella enterica, Alternaria alternata* and *Fusarium oxysporum* (Fig. 2).

**FIG 2.**
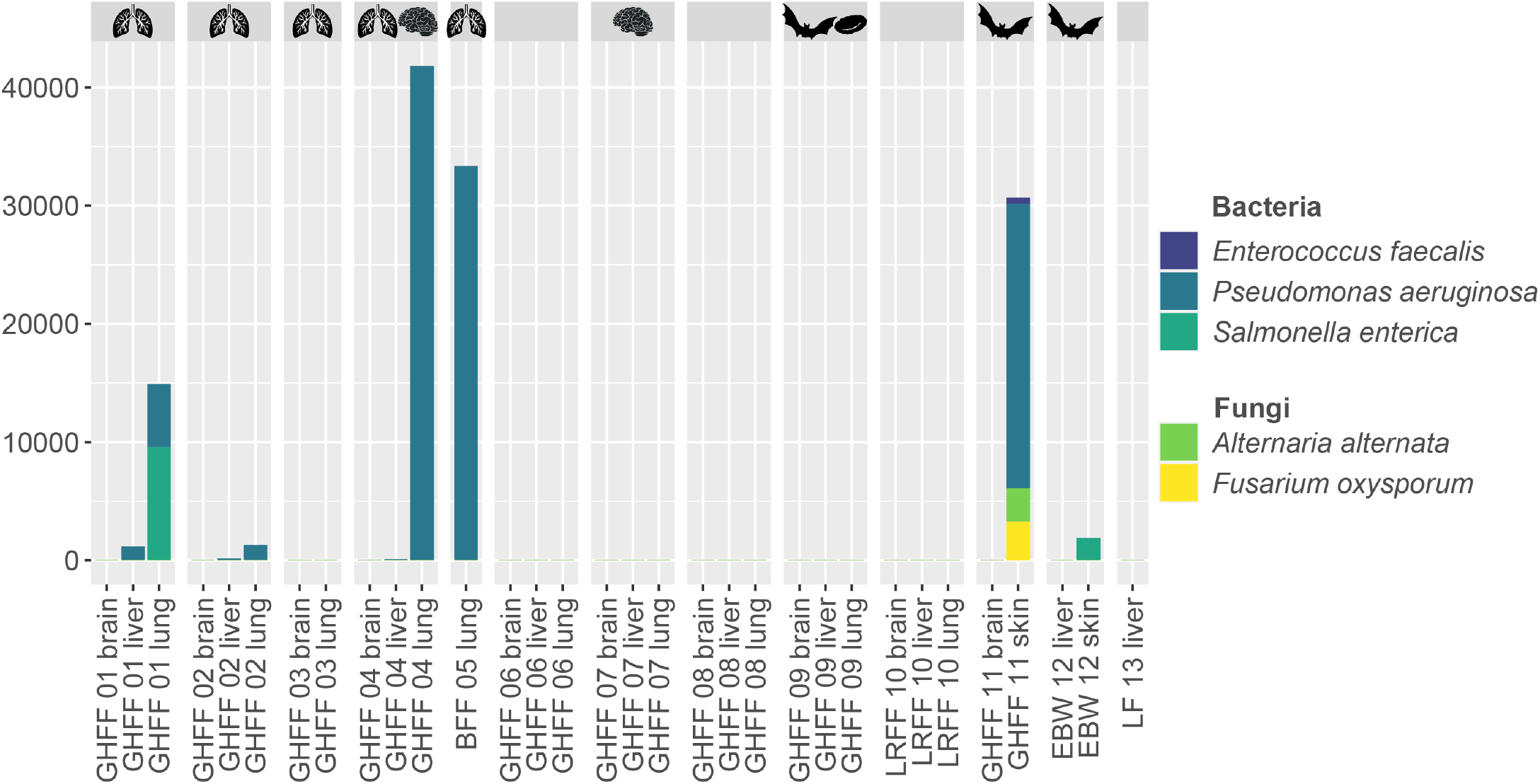
Read abundance displayed as reads per million (RPM) for selected bacterial and fungal families that were chosen based on pathogenic status. Libraries are separated by bat group and the lung, brain, red blood cell and bat silhouettes above the groups indicate which libraries were from bats with lung lesions, neurological signs, leukemia and skin lesions, respectively. Read abundance was calculated using CCMetagen (18).

*P. aeruginosa* (a gram-negative bacterium) had the highest read abundance and was observed in two liver, one brain, one skin and four lung libraries (Figure 2). The four lung libraries were from bats experiencing peracute decline or death with evidence of lung lesions, although we did not detect *P. aeruginosa* in an additional lung library containing bats with similar presentations and one animal with *P. aeruginosa* isolated in lung tissue (group GHFF 03). Notably, the group GHFF 11 skin library, which had comparable *P. aeruginosa* read abundance to the lung libraries from bats with pneumonia, was prepared from skin of three grey-headed flying foxes with noticeable skin lesions (Fig. 2, 1c,d) where *P. aeruginosa* was isolated in culture. The 16S rRNA genes from *P. aeruginosa* from the four positive lung libraries displayed nucleotide sequence identities between 92.5 – 99.4%, with the two most abundant lung libraries, group GHFF 04 and BFF 05, having the most nucleotide sequence identity between the 16S rRNA genes (99.2%). The low sequence identities between the *P. aeruginosa* 16S rRNA genes of groups GHFF01, GHFF02 and GHFF03 should be interpreted with caution as these libraries also had low read coverage, with some sections of the 16S rRNA gene having a coverage of only two reads. Additionally, the *P. aeruginosa* 16S rRNA gene from the skin of group GHFF 11 displayed 97.2% sequence identity to the 16S rRNA sequences from group GHFF 04 and BFF 05. *S. enterica* (gram-negative bacterium) reads were detected in one lung library (group GHFF 01) also containing *P. aeruginosa*, and a skin library from an eastern bent-wing bat (group EBW 12) (Fig. 2). Finally, *E. faecalis* (a gram-positive bacterium), *A. alternata* (fungus) and *F. oxysporum* (fungus) were all observed at lower abundance than *P. aeruginosa* in the group GHFF 11 skin library with skin lesions mentioned above (Fig. 2).

### Overview of the viruses present in bats

Virus contigs matching to 14 families were identified here. Virus contigs belonging to the *Bornaviridae, Adintoviridae, Herpesviridae and Phycodnaviridae* were determined as likely to be endogenous virus elements as they contained no viral conserved domains and/or expected ORFs were interrupted by stop codons. These viral groups were therefore excluded from all analysis. Similarly, those contigs classified in the *Partitiviridae, Tombusviridae, Botourmiaviridae* and *Mitoviridae* were not analysed further as their closest relatives were viruses of plants, fungi and algae suggesting that they are dietary or environmental contaminants. Additionally, a *Picobirnaviridae* contig was disregarded as these viruses likely represent bacteriophage (19). One *Parvoviridae* contig, with the closest BLASTX hit to viruses of the *Dependoparvovirus* genus, was detected in group LF 13 liver at low abundance (Fig. 3), although the sequence was too short (360 bp/ 85 amino acid residues) to perform robust phylogenetic analysis. In contrast, the *Astroviridae, Flaviviridae, Picornaviridae* and *Retroviridae* contigs determined here were considered to be *bona fide* exogenous viruses of vertebrates and therefore investigated further (Fig. 3).

**FIG 3.**
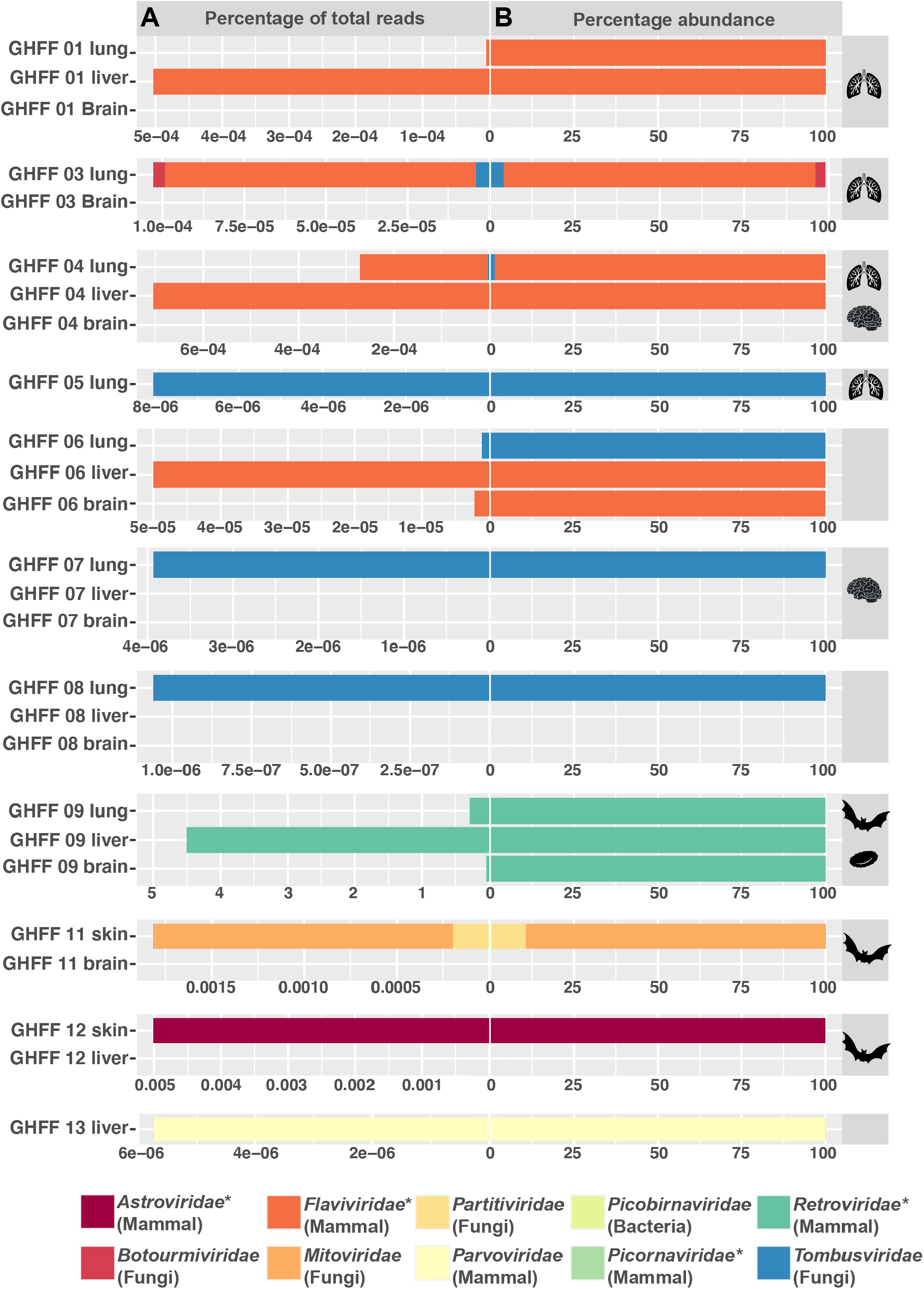
Read abundance of each viral family (excluding viruses determined to be endogenous), presented as (A) the expected count over the total number of trimmed sequence reads for that library multiplied by 100 and (B) the expected count as a percentage of total viral reads for that library. Virus families that are discussed further in this study are highlighted with an asterisk, with the most likely host based on BLASTX for each family shown in parenthesis. Groups with no virus abundance value are not shown. The lung, brain, red blood cell and bat silhouettes above the graphs denote which bat groups had bats with lung lesions, neurological signs, leukemia and skin lesions, respectively.

### Identification of Hervey pteropid gammaretrovirus

A partial retrovirus contig of 8,105 bp was identified in the lung library from group GHFF 09. Sequence comparison over the entire contig revealed a high nucleotide identity (98%) to Hervey pteropid gammaretrovirus (GenBank accession MN413610.1) previously sampled from the faeces of a black flying fox from Queensland, Australia (20). Amino acid sequence identities of the contig discovered here to the gag, pro-pol and env proteins were 100%, 99.2% and 99%, respectively. Such high sequence identities indicate that this represents a variant of Hervey pteropid gammaretrovirus. Complete ORFs were observed for the pro-pol and env genes, although only a partial gag protein missing 22 bp from the 5’ end was recovered (Fig. 4A). The read abundance for Hervey pteropid gammaretrovirus was disproportionately high compared to the other viruses detected here (Fig. 3). The liver library contained the highest read abundance (104,147 FPKM), slightly lower than observed for the host COX1 housekeeping gene (130,674 FPKM). The read abundance for the lung and brain libraries were 51,897 and 4,243 FPKM, respectively. Notably, phylogenetic analysis showed the gammaretroviruses sampled from Australian bats form a clade in the full genome phylogeny, indicative of ongoing evolution within Australia (Fig. 4B).

**FIG 4.**
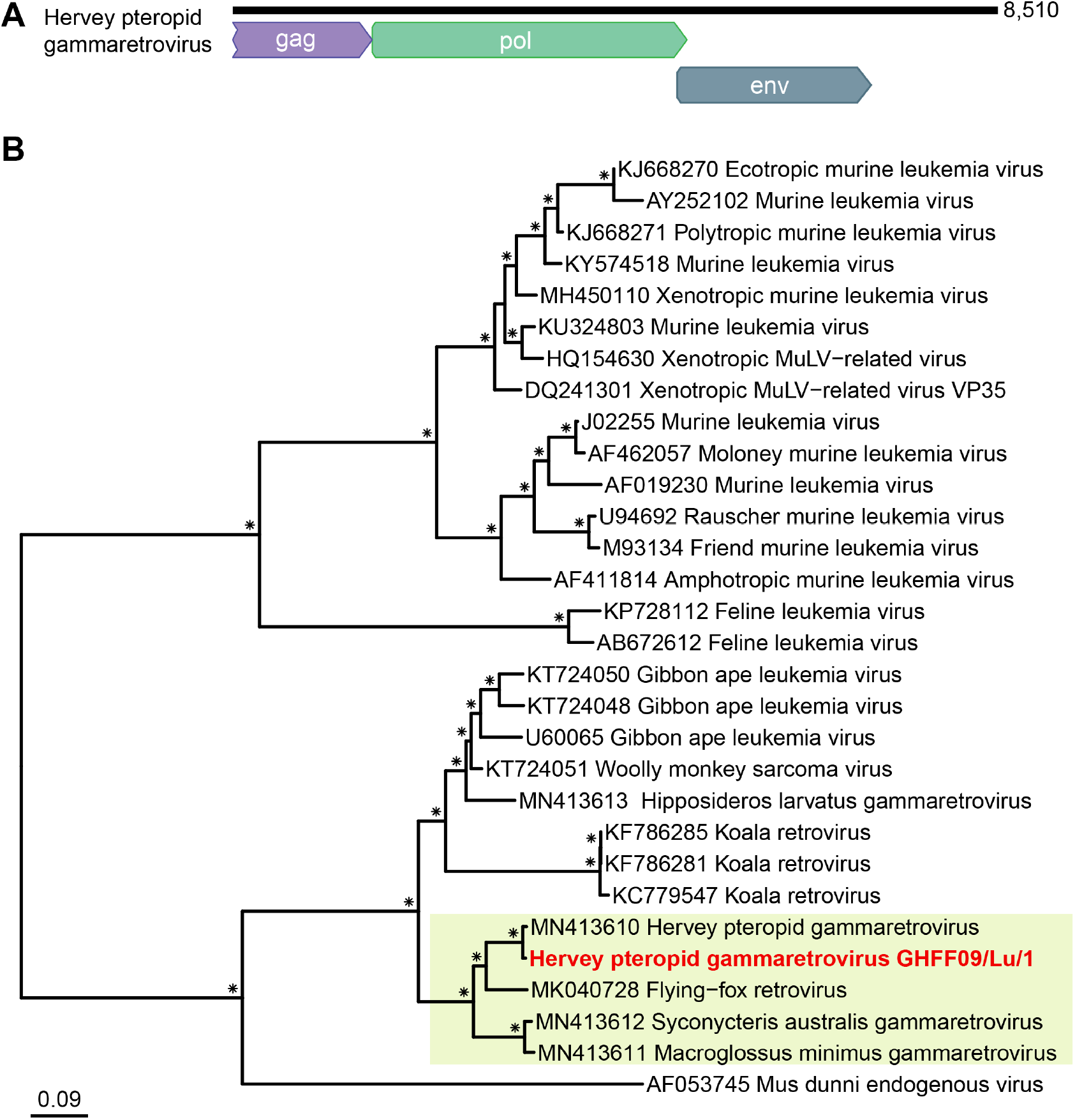
(A) Genome organisation of the Hervey pteropid gammaretrovirus variant detected in this study. (B) Phylogenetic relationships of the gammaretrovirus genus determined using the full genome nucleotide sequence. The bat gammaretrovirus from this study is coloured in red and the gammaretroviruses from Australian bats are highlighted with a yellow background. Bootstrap values >70% are represented by the * symbol shown at the branch node and the tree is rooted at midpoint for clarity. The scale bar represents the number of nucleotide substitutions per site.

We next performed PCR targeting the gag, pol and env genes (Table S2) on the individual lung, brain and liver RNA from the four bats included in group GHFF 09. This revealed that Hervey pteropid gammaretrovirus was present in two grey-headed flying foxes. A positive PCR result was observed in the lung, brain and liver samples from a female grey-headed flying fox found in Sydney, NSW (bat no. 14065.1; Table S1). This bat was euthanised due to ill-thrift and abdominal distension and histological changes were consistent with lymphoid leukemia. An additional positive PCR result was seen for a liver sample from a male grey-headed flying fox with white skin lesions from Woolgoolga, NSW (bat no. 11501.1; Table S1). No RNA from other tissue types was available for PCR for this animal.

### Novel bat astroviruses

We identified a high abundance of astroviruses in a skin library (group EBW 12) from a single male eastern bent-wing bat with noticeable white skin lesions and underlying joint damage associated with severe mite infestation (bat no. 13087.1; Table S1). The eastern bent-wing bat was located in Yass, NSW, a town approximately 300 km from Sydney. Astroviruses possess a positive-sense single strand (+ss) RNA genome and those associated with mammals are classified in the genus *Mamastrovirus* and have been linked to gastroenteritis or neurological issues in some species (21).

Near complete genomes were assembled for three distinct astroviruses, and partial genomes (with at least partial capsid or RdRp) were assembled for a further five distinct astroviruses, with contigs lengths ranging from 6,765 bp to 1443 bp (Fig. 5A). Comparative analysis of the complete capsid protein from the three bat astroviruses with near complete genomes – provisionally denoted bat astrovirus 2 (6,765 bp; 3,377 reads), bat astrovirus 3 (6,748 bp; 3,217 reads) and bat astrovirus 4 (6,747 bp; 2,067 reads) – showed their amino acid identities to each other to be 57-67%, and only 20-55% to other characterised bat astroviruses. Hence, each likely represents a novel virus species based on the current species demarcation in ICTV.

**FIG 5.**
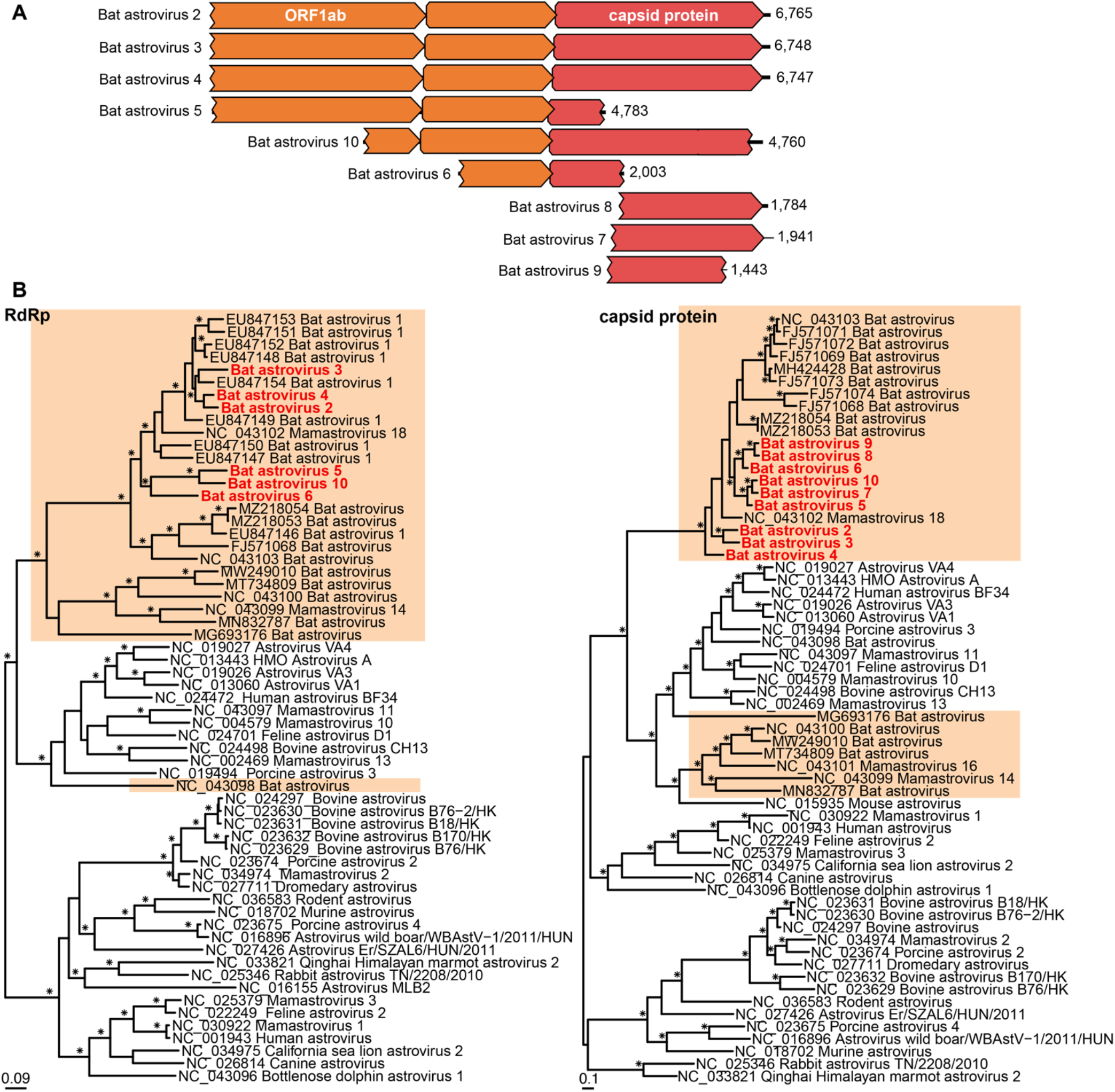
(A) Genomic organisation of the bat astroviruses identified in this study. (B) Phylogenetic relationships of mamastroviruses using the RdRp and capsid protein amino acid sequence. Amino acid alignment lengths were 446 and 465 residues for the RpRp and capsid protein, respectively. Bat astroviruses are highlighted in orange and bat astroviruses from this study are coloured in red. Bootstrap values >70% are represented by the * symbol shown at the branch node. The trees are rooted at midpoint for clarity and the scale bar represents the amino acid substitutions per site.

Phylogenetic analysis of the RdRp and capsid proteins of the bat astrovirus contigs detected here combined with global sequences revealed a clear clustering of astroviruses collected from bats and hence a long-term virus-host association (Fig. 5B). The three new species proposed - bat astrovirus 2, bat astrovirus 3 and bat astrovirus 4 - broadly group together in both the capsid protein and the RdRp phylogenies; although bat astrovirus 3 does not directly cluster with the other two Australian viruses in the RdRp tree due to the inclusion of additional bat astrovirus 1 sequences not present in the capsid tree. Additionally, the partial genome bat astrovirus sequences detected did not group with bat astrovirus 2, 3 and 4, suggesting that multiple lineages of bat astroviruses are evolving in Australia (Fig. 5B).

### Bat pegiviruses

Pegivirus fragments were identified in four bat libraries (group GHFF 01, GHFF 03, GHFF 04 and GHFF 06) containing three, four, five and three bats, respectively (Table S1). Pegiviruses are +ssRNA viruses of the genus *Pegivirus*, family *Flaviviridae*, that are often non-pathogenic in mammalian hosts. In one liver library (group GHFF 04) a complete pegivirus genome (denoted bat pegivirus GHFF04/Li/1) of length 9,784 bp was identified with a read abundance of 1,030, and containing the expected E1, E2, NS2, NS3, NS4A, NS4B, NS5A and NS5B proteins (Fig. 6A). Bat pegivirus GHFF04/Li/1 contigs were also identified in the accompanying lung library from group GHFF 04, with a read abundance of 354. Using primers targeting the NS3 and NS5b region of bat pegivirus GHFF04/Li/1 (Table S2), PCR was performed on the five bats in group GHFF 04. This showed that bat pegivirus GHFF04/Li/1 was present in the lung and liver samples from an adult male grey-headed flying fox from Sydney (bat no. 14121.1; Table S1) that presented with flaccid paralysis and central nervous system depression, including lack of response to stimuli.

**FIG 6.**
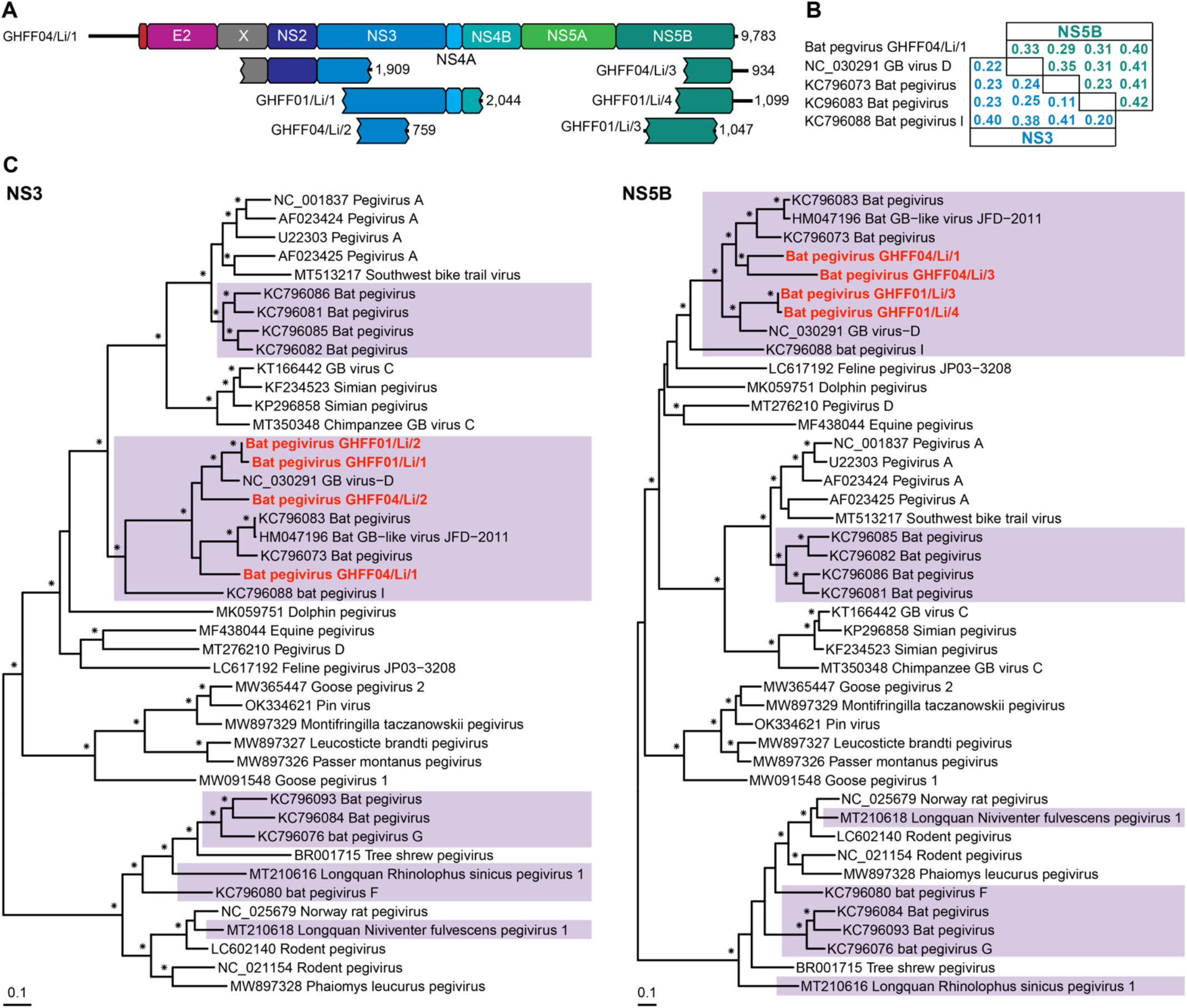
(A) Genomic organisation of the bat pegiviruses identified in this study. (B) Uncorrected (p) distances of the amino acid sequences the NS3 and NS5 proteins of selected bat pegivirus and GHFF04/Li/1. (C) Phylogenetic relationships within the genus *Pegivirus* using the NS3 and NS5B amino acid sequence. Amino acid alignment lengths were 620 residues for the NS3 gene and 531 residues for the NS5B gene. Bat pegiviruses are highlighted in purple and the bat pegiviruses from this study are coloured in red. Bootstrap values >70% are represented by the * symbol shown at the branch node. The tree was midpoint rooted for clarity and the scale bar represents the amino acid substitutions per site.

Necropsy and histopathology revealed necrotising and pyogranulomatous hepatitis, and histiocytic myocarditis. An additional eight unique bat pegivirus contigs (read abundance 221) were identified in liver library GHFF 04, with one contig containing a partial NS3 (GHFF04/Li/2) and another with a partial NS5B (GHFF04/Li/3) region (Fig. 6A). Short pegivirus contigs distinct to the pegiviruses found in GHFF04 were assembled from the liver library from group GHFF 01 (838 reads), the liver library from group GHFF 06 (17 reads), and the lung library from group GHFF 03 (40 reads). As we were unable to extract sufficient RNA from the liver tissue of group GHFF 03, we cannot confirm whether pegivirus contigs were also present in the livers of these bats.

Phylogenetic analysis was conducted on translated contigs of sufficient length (>220 residues) encoding NS3 and NS5b (Fig. 6A). This demonstrated that at least four distinct pegiviruses were present in the sampled bats. The bat pegivirus contigs identified here formed a clade with bat pegiviruses previously identified from species Pegivirus B, suggesting that this species has a long association with bats (Fig. 6C).

### Novel bat kunsagivirus

A novel kunsagivirus (tentatively named Auskunsag virus for the country in which the bat was sampled, <u>Aus</u>tralia) was identified in a single lung and liver library containing three grey-headed flying fox individuals from group GHFF 09. *Kunsagivirus* is a genus of the *Picornaviridae* that have +ssRNA genomes between of 6,800 bp – 7,400 bp in length. The 7,482 bp Auskunsa virus genome comprises the P1 region containing VP0, VP1 and VP3, the P2 region containing the 2A1, 2A2, 2B and 2C and the P3 region containing the 3A, 3B, 3C and 3D (Fig. 7A).

**FIG 7.**
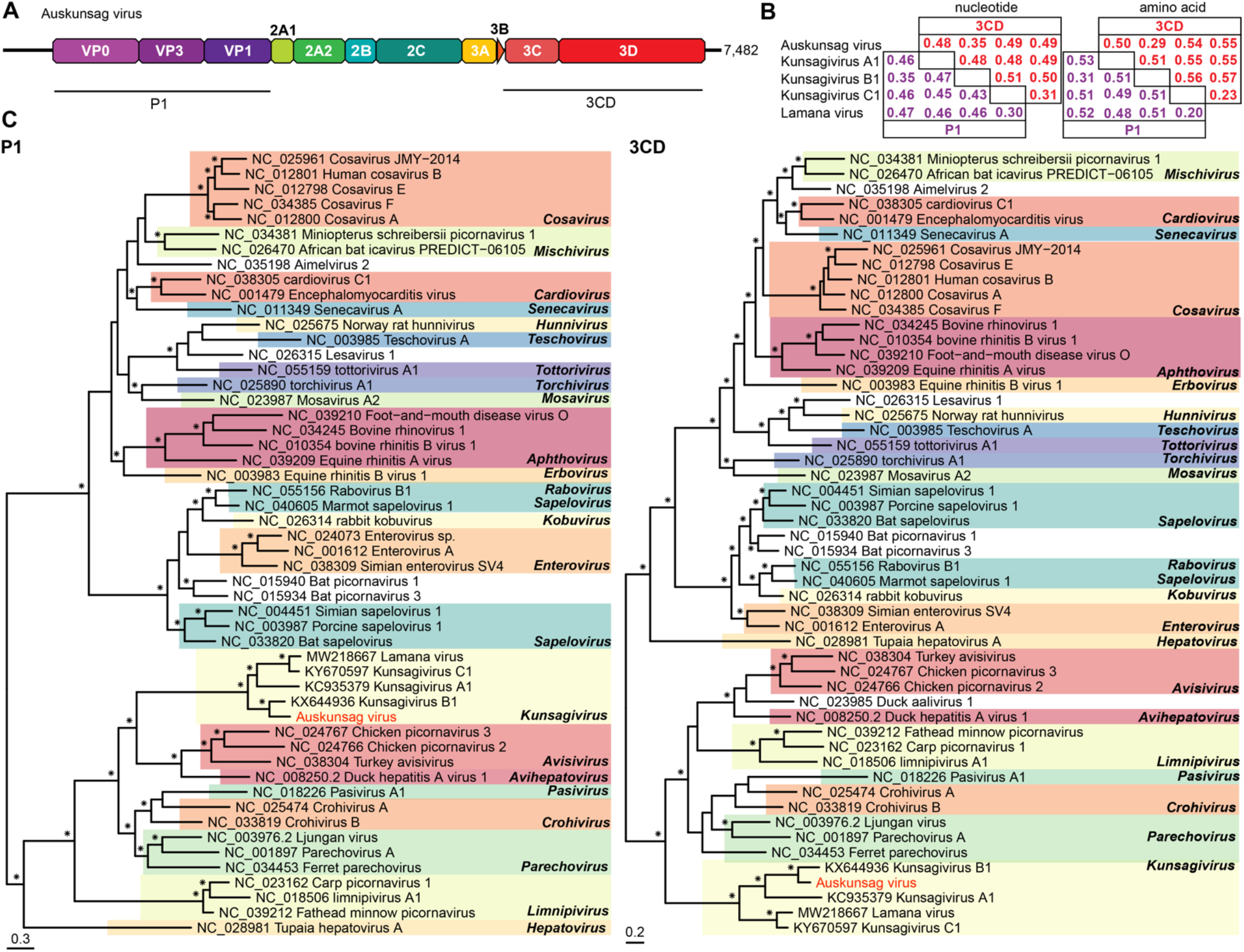
(A) Genomic organisation of the novel kunsagivirus identified in this study. (B) Uncorrected (p) distances among amino acid sequences of the P1 and 3CD regions of the four members of the *Kunsagivirus* genus and Auskunsag virus. (C) Phylogenetic relationships of Auskunsag virus using amino acid sequences of the P1 and 3CD genes. Amino acid alignment lengths were 737 and 653 residues for the P1 and 3CD genes, respectively. Auskunsag virus is coloured in red and the different *Picornaviridae* genera are highlighted in the tree. Bootstrap values >70% are represented by * symbols shown at the branch node. The trees were midpoint rooted for clarity and the scale bar represents the amino acid substitutions per site.

PCR targeting the 3D region (Table S2) was performed on the four individual bats from group GHFF 09, confirming that Auskunsag virus was present the lung and liver sampled from a female juvenile grey-headed flying fox from Sydney with no disease presentation (bat no. 11553.1; Table S1). The read abundance values for the lung and liver libraries were 6,498 and 359, respectively. The closest related virus based on phylogenetic analysis of the P1 and 3CD regions was Kunsagivirus B1 (accession no. KX644936, Fig. 7C) sampled from the faeces of a straw-coloured fruit bat (*Eidolon helvum*) in Cameroon (22). Amino acid identities between Auskunsag virus and Kunsagivirus B1 over the polyprotein, P1 region and 3CD were 68%, 69% and 71%, respectively. The uncorrected (p) distances for the P1 and 3CD region was calculated to determine whether Auskunsag virus should constitute a new species in the *Kunsagivirus* genus. As Auskunsag virus exhibits nucleotide p-distances that fall below the ICTV classification, set at <0.51 for P1 and <0.52 for 3CD, we propose that Auskunsag virus should tentatively represent a new kunsagivirus species (Fig. 7B).

## DISCUSSION

In Australia, numerous native bat species, including the grey-headed flying fox, have experienced population declines that have resulted in their listing as vulnerable or endangered species. Population declines are mainly driven by the effects of climate change and urbanisation, ongoing threats that are likely to continue impacting wild populations (17). Any additional threat to Australian flying foxes, such as infectious disease, could lead population numbers to unrecoverable levels. Given the increasing interest in bat health and the role that bats may play as reservoirs of zoonotic microbes and viruses, we performed metatranscriptomic sequencing on tissue samples from several Australian bats with underlying health issues. From this, we were able to identify several pathogenic bacteria and fungi, an important possibly pathogenic gammaretrovirus, and RNA viruses from the vertebrate-infecting families *Astroviridae, Flaviviridae* and *Picornaviridae*.

A notable aspect of this study was the metatranscriptomic identification of viruses from tissue samples in bats with varied disease syndromes, rather than exploring the healthy state virome. Generally, collecting faeces is the preferred method of sampling bats as it enables large-scale sample collection in a non-invasive manner. Although characterising the faecal virome of bats is important for identifying novel and potentially zoonotic viruses, especially from urban bat populations, the identification of dietary invertebrate and plant viruses is common, such that viromes may differ between faecal and tissue samples (23-25). Our previous study of the faecal virome of the grey-headed flying fox identified bat viruses belonging to the *Coronaviridae* and *Caliciviridae*, as well as a myriad of insect and plant viruses likely associated with the diet (26). Notably, no mammalian viruses belonging to the *Flaviviridae*, as well as only one short contig matching to *Astroviridae* and two short contigs matching to *Picornaviridae*, where identified in marked contrast to the results presented here (26).

Analysis of the microbiome from Australian bat tissues identified four bacterial and two fungal species of pathogenic concern. *Salmonella enterica* serovar Wangata was isolated in culture from the lung and intestine of a young grey-headed flying fox found with histological evidence of colitis, hepatitis and interstitial pneumonia in a pattern consistent with septicaemia (bat no. 13402.1). This organism is an important cause of human salmonellosis in NSW (27). *P. aeruginosa* is an opportunistic pathogen that can cause pneumonia in immunocompromised people (28, 29), which was identified by culture and metatranscriptomic investigation within lung tissue of ten diseased bats in this study.

Affected bats had a consistent pattern of fibrin necrotising, neutrophilic or histiocytic interstitial pneumonia. In two lung libraries, both from flying foxes, *P. aeruginosa* reads were at high abundance (Fig. 2). It is unclear whether the presence of *P. aeruginosa* in these bats is a primary cause of lung disease. All affected animals had a history of trauma, heat stress or involvement in a mass mortality event, suggesting that *P. aeruginosa* is acting as a secondary, opportunistic infection. Furthermore, analysis of the *P. aeruginosa* 16S rRNA gene showed sequence diversity, suggesting that the presentation of peracute pneumonia in the bats is unlikely to be caused by the clonal expansion of a single pathogenic *P. aeruginosa* organism. The abundance of *P. aeruginosa* was also high in a single skin library (Fig. 2).

This library contained ulcerated and hyperkeratotic skin samples from three grey-headed flying foxes where *P. aeruginosa, Serratia marcescens*, and other bacteria were isolated in culture. It is important to note that *P. aeruginosa* is found in the environment and can be found as part of the skin microbiome (28, 30). When *P. aeruginosa* is present on the skin it can opportunistically cause skin and soft tissue infections in humans (30). The high read abundance of *P. aeruginosa* in the skin library of grey-headed flying foxes with noticeable wing skin lesions is interesting, although it may constitute a harmless part of the skin microbiome at the time of sampling.

In some instances, bacteria were isolated in tissue culture of bat lesions, but were not abundant within the metatranscriptomic data, most likely reflecting overgrowth of highly cultivable organisms rather than true organism diversity and abundance. Alternatively, pooling samples for metatranscriptomic investigations may diminish the relative abundance of some organisms. The integration of traditional and metatranscriptomic diagnostic pipelines has the potential to more fully explore the microbial diversity of wildlife while tempering the potential biases inherent in each approach.

A notable finding was the detection of Hervey pteropid gammaretrovirus in a group of grey-headed flying foxes (group GHFF 09). This virus was previously described from the faeces of a black flying fox from Queensland, Australia, and shown to be a functional exogenous virus (20). Investigation herein using PCR revealed that Hervey pteropid gammaretrovirus was present in two grey-headed flying foxes from the same pool of bats. Notably, an adult female grey-headed flying fox with lymphoid leukemia in which virus was detected by PCR in lung, brain and liver samples. The second animal was a male with white skin lesions and detectable virus in the liver. RNA from other tissues was not available for this animal for PCR testing.

These grey-headed flying foxes were from Sydney and Woolgoolga, NSW, with Sydney being the furthest south this virus has been detected (20). Hervey pteropid gammaretrovirus is phylogenetically related to koala retrovirus and gibbon ape leukemia virus (20), both of which are associated with immune deficiencies and leukemia (31, 32). The presence of Hervey pteropid gammaretrovirus in a bat with lymphoid leukemia suggests a possible association with disease, although further research is needed to reveal any mechanistic role the virus plays in disease manifestation. Replication competent Hervey pteropid gammaretrovirus virions have been tested *in vitro* and confirmed to infect bat and human cell lines, although virions were synthetically constructed using the consensus sequence from RNA sequencing (20). Isolating infectious virus from the bat with lymphoid leukemia, combined with additional *in vitro* studies, may provide better insight into transmissibility and the pathogenic potential of the virus. The FPKM counts in the liver library from group GHFF 09 (104,147) were comparable to those for the bat COX1 housekeeping gene (130,674). Such a high abundance is compatible with active virus replication at the time of sampling, although the contribution of each individual bat liver sample toward the total liver library gammaretrovirus abundance cannot be determined.

In the same group of bats (GHFF 09) in which we identified Hervey pteropid gammaretrovirus we detected a novel picornavirus belonging to the genus *Kunsagivirus*. This genus currently contains three recognised species sampled from the faeces of a European roller in 2011 (33) and a straw-coloured fruit bat in Cameroon (22), respectively, and from the blood of a yellow baboon in Tanzania (34). A fourth kunsagivirus sequence was more recently sampled from vervet monkeys from Uganda (35). Here, we characterised a novel kunsagivirus, tentatively named Auskunsag virus, in a liver sample from a female juvenile grey-headed flying fox from Sydney, Australia, and absent from the two bats in which Hervey pteropid gammaretrovirus was detected. Auskunsag virus is the first report of a kunsagivirus in the Asia-pacific region and of a kunsagivirus in tissue samples, indicating that this genus is most likely mammalian-infecting and not dietary or invertebrate-associated. Viruses of the genus *Kunsagivirus* currently have no disease association, although additional research is needed for confirmation.

Multiple astroviruses were detected in a single skin library containing one eastern-bent wing bat that had noticeable skin lesions coupled with underlying join damage. A total of 74% of the characterised bat astroviruses on NCBI GenBank were sampled exclusively from faeces. As it currently stands, no bat astroviruses have been sampled from skin, although Avian nephritis virus 3 of the genus *Avastrovirus* has been detected in the joint and tendons sampled from boiler chickens and poult turkeys with arthritis and tenosynovitis and was proposed to have a possible association with these conditions (36). The diversity of astroviruses in Australian bats has yet to be assessed, and only one sequence of the *Astroviridae* from an Australian bat is available on GenBank. This sequence, microbat bastrovirus (accession no. MT766313), is more closely related to the diverse group of astroviruses termed bastroviruses that contain a hepe-like non-structural protein and an astro-like structural protein (37). A study of Asian and European bat species showed that astroviruses are common in bat populations and in some incidences were at high prevalence (38-42). The detection of several diverse bat astroviruses in this study suggests that numerous astroviruses may be circulating within the bat population in Australia and that further research is needed to fully understand their community structure and potential contribution to disease.

Finally, we detected bat pegivirus contigs in four grey-headed flying fox groups (GHFF 01, GHFF 03, GHFF 04 and GHFF 06), with one containing a complete genome – bat pegivirus GHFF04/Li/1. Generally, pegiviruses are non-pathogenic and associated with persistent infections in mammals, although members of the species Pegivirus D have been associated with the development of Theiler’s disease in infected horses (43). Although we detected bat pegivirus GHFF04/Li/1 in a grey-headed flying fox with necrotising and granulomatous hepatitis and histiocytic myocarditis, we also detected bat pegiviruses in bats with no clear liver or heart disease suggesting that it is likely a commensal pathogen like most other pegiviruses.

The sustainability of micro and megabat species globally is important for the sustainably of the world’s ecosystems. In countries with continual threats, such as habitat change and destruction, land use change, agricultural practices adverse to bat health, extreme temperatures and white-nose syndrome, efforts to maintain population numbers are critical. Ongoing monitoring of bat health and disease and using traditional and metatranscriptomic diagnostic techniques is warranted to explore the diversity of microbes with pathogenic potential that might be expressed in the form of disease in populations subject to landscape-wide change. The presence of a retrovirus from a genus with members that are associated with immune deficiencies and leukemia, in a bat with lymphoid leukemia, undoubtedly merits further investigation and broader surveillance.

## METHODS

### Animal Ethics and sample collection

Wild bats were examined under a License to Rehabilitate Injured, Sick or Orphaned Protected Wildlife (no. MWL000100542) issued by the NSW Department of the Environment All samples were collected post-mortem from bats that were submitted for disease investigation. These samples were collected under the auspices of the Taronga Conservation Society Australia’s Animal Ethics Committee (approval no. 3b1218), pursuant to NSW Office of Environment and Heritage-issued scientific license no. SL10469 and SL100104.

Samples from 38 grey-headed flying foxes, one black flying fox, one large footed myotis, one eastern bent-wing bat and two little red flying foxes were collected between November 2013 and April 2021 (Table S1). Most animals emanated from the Sydney basin and the central coast region of New South Wales. Fresh portions of brain, lung, liver, skin, heart, kidney and any lesions were collected aseptically post-mortem and frozen at –80°C. Impression smears of cut sections of lung tissue, or other lesions, were prepared from a subset of animals. The tissue was blotted onto a glass slide, air dried, fixed and stained with Quick Dip (Fronine, Thermo Fisher Scientific Aust. Pty. Ltd., Scoresby, Victoria, Australia) and examined at 1000x magnification with oil emersion microscopy. A range of tissues from each animal was fixed in 10% neutral buffered formalin, processed in ethanol, embedded with paraffin, sectioned, stained with hematoxylin and eosin, and mounted with a cover slip for examination by light microscopy.

### Microbial culture

Microbial culture was conducted using a subset of bat tissues where there were gross or histological lesions, including 16 lung samples, and four skin samples. Culture was also conducted on four food sources used in bat rehabilitation care. Lung and lesion impression smears were stained with Gram and ZN were examined under 1000x oil immersion microscopy. Skin lesions were swabbed with a sterile applicator and the pleural surface of each lung sample was seared with a hot scalpel blade, and a sterile microbial loop was inserted into the deeper tissue to inoculate horse blood agar (HBA) and MacConkey agar (MAC) (Thermo Fisher Scientific, Scoresby, Victoria, Australia), which were incubated at 35°C in 4.5% carbon dioxide for 24–48 h. Isolates were identified with API 20 NE identification kits (bioMerieux SA, Lyon, France) using manufacturer’s instructions.

### Sample preparation, library construction and virus discovery

Tissue samples were homogenised using 2.38 mm metal beads (Qiagen) in the TissueLyser LT (Qiagen). Total RNA was extracted using the RNeasy Plus Mini Kit (Qiagen) following the manufacturer’s protocol. Extracted RNA was pooled according to the syndrome, tissue type and bat species (Table S1). Sequencing libraries were constructed using the Illumina Stranded Total RNA Prep with Ribo-Zero Plus (Illumina) preparation kit after the removal of rRNA. Libraries were sequenced on the Illumina NovaSeq platform as 150 bp paired end reads at the Australian Genome Research Facility (AGRF, Melbourne). Sequencing reads that contained read ends with a phred score below 25 and adapter sequences were quality trimmed using cutadapt version 1.8.3 (44). Trimmed reads were then *de novo* assembled into contigs using Megahit version 1.1.3 (45, 46). The resulting contigs were then compared to the non-redundant protein database using Diamond version 2.0.9 with an e-value cut-off of 1E^-5^. Hervey pteropid gammaretrovirus was identified using an in-house retrovirus discovery pipeline (W.S Chang, A.S Baez-Ortiz, J.E Mahar, E.C Holmes, C Le Lay, K Rose and C Blaker, manuscript in preparation). Attempts to extend virus contigs were made by using a reassembling megahit contigs using a Geneious version 2022.1.1 assembler.

### Taxonomy profiling and abundance calculation

Taxonomic assignment and abundance information for bacterial, fungal and metazoan contigs was accessed using CCMetagen version 1.2.4 (18) and kma version 1.2.4 (47). Taxonomic information for viral contigs was retrieved from the protein database results. Read abundance values for the virus contigs and COX1 genes (accession no. KF726143 for *Pteropus* species, MK410364 for the eastern bent-wing bat and a COX1 contig identified in the large-footed myotis library) were calculated by mapping trimmed sequencing reads using the RSEM version 1.3.2 tool (48) in Trinity and Bowtie2 version 2.3.3.1 (49, 50). The 16S rRNA genes from the *P. aeruginosa* in the bat lung and skin libraries were obtained by mapping trimmed reads to a *P. aeruginosa* 16S rRNA gene available on NCBI GenBank (accession no. CP003149) and extracting the 0% majority consensus (i.e., least ambiguities in sequence) using Geneious version 2022.1.1.

### Phylogenetic analysis

Amino acid and nucleotide alignments of virus genes were generated using MAFFT version 7.450 and the E-INS-I algorithm (51), with ambiguously aligned regions removed using the gappyout method in TrimAL version 1.4.1 (52). The model finder program (53) in IQ-TREE version 1.6.7 (54) was used to determine the best-fit models of amino acid and nucleotide substitution and the same program was used to infer maximum likelihood trees. Ultrafast bootstrapping with 1000 replicates was to provide an indication of nodal support, and the nearest neighbour interchange was applied to search for optimal tree structure (55).

## Data availability

The sequencing data for this study are available on the NCBI Sequence Read Archive database under the BioProject no. PRJNA885898 and SRA no. SRR21780604 – SRR21780623. The virus genomes have been deposited on NCBI GenBank under the accession no. OP589976-OP589993.

## ACKNOWLEDGEMENTS

We thank Natalie Miller and Paul Thompson, Taronga Wildlife Hospital, for the microbiological examination of tissue samples. We are grateful for the care and expertise exhibited by those responding to injured bats, including Meg Churches, Storm Stanford, Sarah Curran - Wildlife Information, Rescue, and Education Service (WIRES), Dr. Kerryn Parry-Jones – Wildlife Animal Rescue and Care, Dr. Stephen Deist – Pacific Vet Care and Mid-North Coast WIRES, Erin Stokes and Dr. Amber Brett - North Nowra Vet Hospital and Wildlife Rescue South Coast, Dr. Louise Hayles - Kippax Veterinary Hospital, Denise Kay – ACT Wildlife, Sandra Guy – Sydney Wildlife Rescue, and Drs. John Martin, and Phil Kemsley. Taronga Conservation Society Australia is acknowledged for ongoing funding of the Australian Registry of Wildlife Health. This project received funding from the Australian Government’s Wildlife Rescue and Rehabilitation initiative, and by an Australian Research Council Australian Laureate Fellowship to ECH (FL170100022).

## Notes

### Competing Interest Statement

The authors have declared no competing interest.

## REFERENCES

1. Mollentze N, Streicker DG. 2020. Viral zoonotic risk is homogenous among taxonomic orders of mammalian and avian reservoir hosts. Proc Natl Acad Sci USA 117:9423–9430.

2. Zhou H, Ji J, Chen X, Bi Y, Li J, Wang Q, Hu T, Song H, Zhao R, Chen Y, Cui M, Zhang Y, Hughes AC, Holmes EC, Shi W. 2021. Identification of novel bat coronaviruses sheds light on the evolutionary origins of SARS-CoV-2 and related viruses. Cell 184:4380-4391.e14.

3. Wu Z, Yang L, Ren X, He G, Zhang J, Yang J, Qian Z, Dong J, Sun L, Zhu Y, Du J, Yang F, Zhang S, Jin Q. 2016. Deciphering the bat virome catalog to better understand the ecological diversity of bat viruses and the bat origin of emerging infectious diseases. ISME J 10:609–20.

4. Wang J, Anderson DE, Halpin K, Hong X, Chen H, Walker S, Valdeter S, van der Heide B, Neave MJ, Bingham J, O’Brien D, Eagles D, Wang LF, Williams DT. 2021. A new Hendra virus genotype found in Australian flying foxes. Virol J 18:197.

5. Mishra N, Fagbo SF, Alagaili AN, Nitido A, Williams SH, Ng J, Lee B, Durosinlorun A, Garcia JA, Jain K, Kapoor V, Epstein JH, Briese T, Memish ZA, Olival KJ, Lipkin WI. 2019. A viral metagenomic survey identifies known and novel mammalian viruses in bats from Saudi Arabia. PLoS One 14:e0214227.

6. Irving AT, Ahn M, Goh G, Anderson DE, Wang LF. 2021. Lessons from the host defences of bats, a unique viral reservoir. Nature 589:363–370.

7. Banerjee A, Baker ML, Kulcsar K, Misra V, Plowright R, Mossman K. 2020. Novel insights into immune systems of bats. Front Immunol 11:26.

8. Hoyt JR, Kilpatrick AM, Langwig KE. 2021. Ecology and impacts of white-nose syndrome on bats. Nat Rev Microbiol 19:196–210.

9. Cheng TL, Reichard JD, Coleman JTH, Weller TJ, Thogmartin WE, Reichert BE, Bennett AB, Broders HG, Campbell J, Etchison K, Feller DJ, Geboy R, Hemberger T, Herzog C, Hicks AC, Houghton S, Humber J, Kath JA, King RA, Loeb SC, Massé A, Morris KM, Niederriter H, Nordquist G, Perry RW, Reynolds RJ, Sasse DB, Scafini MR, Stark RC, Stihler CW, Thomas SC, Turner GG, Webb S, Westrich BJ, Frick WF. 2021. The scope and severity of white-nose syndrome on hibernating bats in North America. Conserv Biol 35:1586–1597.

10. Banyard AC, Hayman D, Johnson N, McElhinney L, Fooks AR. 2011. Bats and lyssaviruses. Adv Virus Res 79:239–89.

11. International Union for Consevation of Nature. 2021. The IUCN Red List of Threatened Species. https://www.iucnredlist.org.

12. International Union for Conservation of Nature. 2021. Data from “Pteropus poliocephalus”. The IUCN Red List of Threatened Species https://www.iucnredlist.org/species/18751/22085511.

13. Kolkert H, Andrew R, Smith R, Rader R, Reid N. 2020. Insectivorous bats selectively source moths and eat mostly pest insects on dryland and irrigated cotton farms. Ecol Evol 10:371–388.

14. Law BS, Lean M. 1999. Common blossom bats (Syconycteris australis) as pollinators in fragmented Australian tropical rainforest. Biol Conserv 91:201–212.

15. Moran C, Catterall CP, Kanowski J. 2009. Reduced dispersal of native plant species as a consequence of the reduced abundance of frugivore species in fragmented rainforest. Biol conserv 142:541–552.

16. Mo M, Roache M, Davies J, Hopper J, Pitty H, Foster N, Guy S, Parry-Jones K, Francis G, Koosmen A. 2021. Estimating flying-fox mortality associated with abandonments of pups and extreme heat events during the austral summer of 2019–20. Pacific Conserv Biol 28:124–139.

17. Welbergen JA, Klose SM, Markus N, Eby P. 2008. Climate change and the effects of temperature extremes on Australian flying-foxes. Proc Biol Sci 275:419–25.

18. Marcelino VR, Clausen P, Buchmann JP, Wille M, Iredell JR, Meyer W, Lund O, Sorrell TC, Holmes EC. 2020. CCMetagen: comprehensive and accurate identification of eukaryotes and prokaryotes in metagenomic data. Genome Biol 21:103.

19. Krishnamurthy SR, Wang D. 2018. Extensive conservation of prokaryotic ribosomal binding sites in known and novel picobirnaviruses. Virology 516:108–114.

20. Hayward JA, Tachedjian M, Kohl C, Johnson A, Dearnley M, Jesaveluk B, Langer C, Solymosi PD, Hille G, Nitsche A, Sánchez CA, Werner A, Kontos D, Crameri G, Marsh GA, Baker ML, Poumbourios P, Drummer HE, Holmes EC, Wang LF, Smith I, Tachedjian G. 2020. Infectious KoRV-related retroviruses circulating in Australian bats. Proc Natl Acad Sci USA 117:9529–9536.

21. De Benedictis P, Schultz-Cherry S, Burnham A, Cattoli G. 2011. Astrovirus infections in humans and animals - molecular biology, genetic diversity, and interspecies transmissions. Infect Genet Evol 11:1529–44.

22. Yinda CK, Zeller M, Conceição-Neto N, Maes P, Deboutte W, Beller L, Heylen E, Ghogomu SM, Van Ranst M, Matthijnssens J. 2016. Novel highly divergent reassortant bat rotaviruses in Cameroon, without evidence of zoonosis. Sci Rep 6:34209.

23. Li L, Victoria JG, Wang C, Jones M, Fellers GM, Kunz TH, Delwart E. 2010. Bat guano virome: predominance of dietary viruses from insects and plants plus novel mammalian viruses. J Virol 84:6955–65.

24. Cobbin JC, Charon J, Harvey E, Holmes EC, Mahar JE. 2021. Current challenges to virus discovery by meta-transcriptomics. Curr Opin Virol 51:48–55.

25. Hardmeier I, Aeberhard N, Qi W, Schoenbaechler K, Kraettli H, Hatt JM, Fraefel C, Kubacki J. 2021. Metagenomic analysis of fecal and tissue samples from 18 endemic bat species in Switzerland revealed a diverse virus composition including potentially zoonotic viruses. PLoS One 16:e0252534.

26. Van Brussel K, Mahar JE, Ortiz-Baez AS, Carrai M, Spielman D, Boardman WSJ, Baker ML, Beatty JA, Geoghegan JL, Barrs VR, Holmes EC. 2022. Faecal virome of the Australian grey-headed flying fox from urban/suburban environments contains novel coronaviruses, retroviruses and sapoviruses. Virology 576:42–51.

27. Simpson KMJ, Mor SM, Ward MP, Walsh MG. 2019. Divergent geography of Salmonella wangata and Salmonella typhimurium epidemiology in New South Wales, Australia. One Health 7:100092.

28. Moradali MF, Ghods S, Rehm BH. 2017. Pseudomonas aeruginosa lifestyle: a paradigm for adaptation, survival, and persistence. Front Cell Infect Microbiol 7:39.

29. Reynolds D, Kollef M. 2021. The epidemiology and pathogenesis and treatment of Pseudomonas aeruginosa infections: an Update. Drugs 81:2117–2131.

30. Spernovasilis N, Psichogiou M, Poulakou G. 2021. Skin manifestations of Pseudomonas aeruginosa infections. Curr Opin Infect Dis 34:72–79.

31. Xu W, Eiden MV. 2015. Koala retroviruses: evolution and disease dynamics. Annu Rev Virol 2:119–34.

32. Kawakami TG, Huff SD, Buckley PM, Dungworth DL, Synder SP, Gilden RV. 1972. C-type virus associated with gibbon lymphosarcoma. Nat New Biol 235:170–1.

33. Boros Á, Kiss T, Kiss O, Pankovics P, Kapusinszky B, Delwart E, Reuter G. 2013. Genetic characterization of a novel picornavirus distantly related to the marine mammal-infecting aquamaviruses in a long-distance migrant bird species, European roller (Coracias garrulus). J Gen Virol 94:2029–2035.

34. Buechler CR, Bailey AL, Lauck M, Heffron A, Johnson JC, Campos Lawson C, Rogers J, Kuhn JH, O’Connor DH. 2017. Genome sequence of a novel Kunsagivirus (Picornaviridae: Kunsagivirus) from a wild baboon (Papio cynocephalus). Genome Announc 5:e00261–17.

35. Kuhn JH, Sibley SD, Chapman CA, Knowles NJ, Lauck M, Johnson JC, Lawson CC, Lackemeyer MG, Valenta K, Omeja P, Jahrling PB, O’Connor DH, Goldberg TL. 2020. Discovery of Lanama virus, a distinct member of species Kunsagivirus C (Picornavirales: Picornaviridae), in wild vervet monkeys (Chlorocebus pygerythrus). Viruses 12:1436.

36. de Wit JJ, Dam GB, de Laar JM, Biermann Y, Verstegen I, Edens F, Schrier CC. 2011. Detection and characterization of a new astrovirus in chicken and turkeys with enteric and locomotion disorders. Avian Pathol 40:453–61.

37. Oude Munnink BB, Cotten M, Canuti M, Deijs M, Jebbink MF, van Hemert FJ, Phan MV, Bakker M, Jazaeri Farsani SM, Kellam P, van der Hoek L. 2016. A novel astrovirus-like RNA virus detected in human stool. Virus Evol 2:vew005.

38. Lacroix A, Duong V, Hul V, San S, Davun H, Omaliss K, Chea S, Hassanin A, Theppangna W, Silithammavong S, Khammavong K, Singhalath S, Afelt A, Greatorex Z, Fine AE, Goldstein T, Olson S, Joly DO, Keatts L, Dussart P, Frutos R, Buchy P. 2017. Diversity of bat astroviruses in Lao PDR and Cambodia. Infect Genet Evol 47:41–50.

39. Fischer K, Zeus V, Kwasnitschka L, Kerth G, Haase M, Groschup MH, Balkema-Buschmann A. 2016. Insectivorous bats carry host specific astroviruses and coronaviruses across different regions in Germany. Infect Genet Evol 37:108–16.

40. Kemenesi G, Dallos B, Görföl T, Boldogh S, Estók P, Kurucz K, Kutas A, Földes F, Oldal M, Németh V, Martella V, Bányai K, Jakab F. 2014. Molecular survey of RNA viruses in Hungarian bats: discovering novel astroviruses, coronaviruses, and caliciviruses. Vector Borne Zoonotic Dis 14:846–55.

41. Zhu HC, Chu DKW, Liu W, Dong BQ, Zhang SY, Zhang JX, Li LF, Vijaykrishna D, Smith GJD, Chen HL, Poon LLM, Peiris JSM, Guan Y. 2009. Detection of diverse astroviruses from bats in China. J Gen Virol 90:883–887.

42. Fischer K, Pinho Dos Reis V, Balkema-Buschmann A. 2017. Bat astroviruses: towards understanding the transmission dynamics of a neglected virus family. Viruses 9:34.

43. Chandriani S, Skewes-Cox P, Zhong W, Ganem DE, Divers TJ, Van Blaricum AJ, Tennant BC, Kistler AL. 2013. Identification of a previously undescribed divergent virus from the Flaviviridae family in an outbreak of equine serum hepatitis. Proc Natl Acad Sci USA 110:E1407–15.

44. Kechin A, Boyarskikh U, Kel A, Filipenko M. 2017. cutPrimers: a new tool for accurate cutting of primers from reads of targeted next generation sequencing. J Comput Biol 24:1138–1143.

45. Li D, Liu CM, Luo R, Sadakane K, Lam TW. 2015. MEGAHIT: an ultra-fast single-node solution for large and complex metagenomics assembly via succinct de Bruijn graph. Bioinformatics 31:1674–6.

46. Li D, Luo R, Liu CM, Leung CM, Ting HF, Sadakane K, Yamashita H, Lam TW. 2016. MEGAHIT v1.0: A fast and scalable metagenome assembler driven by advanced methodologies and community practices. Methods 102:3–11.

47. Clausen P, Aarestrup FM, Lund O. 2018. Rapid and precise alignment of raw reads against redundant databases with KMA. BMC Bioinformatics 19:307.

48. Li B, Ruotti V, Stewart RM, Thomson JA, Dewey CN. 2010. RNA-Seq gene expression estimation with read mapping uncertainty. Bioinformatics 26:493–500.

49. Langmead B, Wilks C, Antonescu V, Charles R. 2019. Scaling read aligners to hundreds of threads on general-purpose processors. Bioinformatics 35:421–432.

50. Langmead B, Salzberg SL. 2012. Fast gapped-read alignment with Bowtie 2. Nat Methods 9:357–9.

51. Katoh K, Standley DM. 2013. MAFFT multiple sequence alignment software version 7: improvements in performance and usability. Mol Biol Evol 30:772–80.

52. Capella-Gutiérrez S, Silla-Martínez JM, Gabaldón T. 2009. trimAl: a tool for automated alignment trimming in large-scale phylogenetic analyses. Bioinformatics 25:1972–3.

53. Kalyaanamoorthy S, Minh BQ, Wong TKF, von Haeseler A, Jermiin LS. 2017. ModelFinder: fast model selection for accurate phylogenetic estimates. Nat Methods 14:587–589.

54. Nguyen L-T, Schmidt HA, von Haeseler A, Minh BQ. 2014. IQ-TREE: A fast and effective stochastic algorithm for estimating maximum-likelihood phylogenies. Mol Biol Evol 32:268–274.

55. Hoang DT, Chernomor O, von Haeseler A, Minh BQ, Vinh LS. 2017. UFBoot2: Improving the ultrafast bootstrap approximation. Mol Biol Evol 35:518–522.

