## Supplementary Information for "Gammaretroviruses, novel viruses and pathogenic bacteria in Australian bats with neurological signs, pneumonia and skin lesions"

**SUPPLEMENTARY MATERIAL**

**Supplementary Table 1.** Overview of tissue type included in each library pool for each individual bat, species, sex, age, location of sampling and disease presentation observed.

| Group | Bat no. | Species | Sex | Age | Sample location | Tissue included in library | | | | Clinical signs and histopathology |
|---|---|---|---|---|---|---|---|---|---|---|
| GHFF 01 | 9599.2 | *Pteropus poliocephalus* (Grey-headed flying fox) | | Juvenile | | NA | Brain | Liver | NA | MM, NIP |
| | 11573.4 | | Male | Juvenile | Centennial Park | Lung | Brain | Liver | NA | MM, NIP |
| | 9599.9 | | | Juvenile | | Lung | Brain | Liver | NA | MM, NIP |
| | 13402.1 | | Female | Juvenile | Wyoming | Lung | Brain | NA | NA | MM, NIP |
| GHFF 02 | 14054.1 | | | | | NA | Brain | Liver | NA | Pleuropneumonia |
| | 13906.1 | | | | | Lung | NA | Liver | NA | NIP, HP |
| | 13940.1 | | Female | Subadult | Centennial Park | Lung | Brain | Liver | NA | NIP |
| | 13977.1 | | | | | Lung | Brain | Liver | NA | NIP |
| | 14052.1 | | | | | Lung | Brain | NA | NA | NIP |
| GHFF 03 | 14093.4 | | | | | Lung | NA | NA | NA | NIP |
| | 10124.1 | | Male | Adult | Marsfield | Lung | NA | NA | NA | NIP |
| | 12126.1 | | Male | Juvenile | Clovelly | Lung | NA | NA | NA | NIP |
| | 14003.1 | | | | | Lung | Brain | NA | NA | NIP |
| | 13955.1 | | | | | NA | Brain | NA | NA | NIP |
| | 13997.1 | | | | | NA | Brain | NA | NA | NIP |
| GHFF 04 | 14058.1 | | | | | Lung | Brain | Liver | NA | HP, neurological |
| | 14053.1 | | | | | Lung | Brain | Liver | NA | NIP, neurological |
| | 14088.1 | | | Adult | | Lung | Brain | Liver | NA | NIP, HP, neurological |
| | 14093.2 | | | | | NA | Brain | Liver | NA | NIP, neurological |
| | 14121.1 | | Male | Adult | Kirribilli | NA | NA | Liver | NA | NIP, neurological, granulomatous |
| GHFF 06 | 11573.1 | | Female | Juvenile | Centennial Park | Lung | Brain | Liver | NA | MM |
| | 11573.2 | | | Juvenile | | Lung | Brain | NA | NA | MM |
| | 13368.1 | | | Juvenile | | NA | NA | Liver | NA | MM |
| | 13402.2 | | Female | Juvenile | Wyoming | NA | NA | Liver | NA | MM, NIP |

| | | | | | | | | | | |
|---|---|---|---|---|---|---|---|---|---|---|
| GHFF 07 | 14094.4 | | | | | Lung | Brain | Liver | NA | Neurological |
| | 14123.1 | | Male | Adult | Lismore | Lung | Brain | Liver | NA | Paralysis |
| | 14089.1 | | Male | Adult | Kangaroo Valley | Lung | NA | NA | NA | Neurological |
| GHFF 08 | 13998.1 | | | | | Lung | Brain | NA | NA | |
| | 14120.1 | | Female | Subadult | Kingsford | Lung | Brain | Liver | NA | |
| | 13471.1 | | | | | NA | Brain | NA | NA | |
| | 14048.1 | | | | | Lung | Brain | NA | NA | Trauma |
| GHFF 09 | 14065.1 | | Female | Adult | Sydney | Lung | Brain | Liver | NA | Lymphoid leukemia |
| | 11553.1 | | Female | Juvenile | Botanic Gardens | Lung | Brain | Liver | NA | Euthanised |
| | 11501.1 | | Male | Subadult | Woolgoolga | NA | NA | Liver | NA | WSL, mites |
| | 13905.1 | | | | | NA | Brain | NA | NA | |
| GHFF 11 | 14130.1 | | Female | Subadult | | NA | Brain | NA | Skin | WSL |
| | 14130.3 | | Female | Juvenile | | NA | Brain | NA | Skin | WSL |
| | 14130.2 | | Female | Juvenile | | NA | Brain | NA | Skin | WSL |
| BFF 05 | 13961.1 | *Pteropus alecto* (Black flying fox) | | | | Lung | NA | NA | NA | NIP, neurological |
| LRFF 10 | 14064.1 | Pteropus *scapulatus* (Little red flying fox) | Female | | | Lung | Brain | Liver | NA | Trauma |
| | 14064.2 | | | Foetus | | NA | Brain | Liver | NA | Trauma |
| EBW 12 | 13087.1 | *Miniopterus orianae oceanensis* (Eastern bent-wing bat) | Male | Subadult | Yass | NA | NA | Liver | Skin | WSL |
| LF 13 | 10628.1 | *Myotis Macropus* (Large footed myotis) | Male | Adult | Castle Hill | NA | NA | Liver | NA | Predation |

[orange] = Australian Capital Territory, [blue] = New South Wales
NA = Not available/not included, Lung, Brain, Skin = Diseased tissue included in library, [yellow] = Control
MM = Mass mortality, NIP = Neutrophilic interstitial pneumonia (lung lesions), HP = Histiocytic pneumonia (lung lesions), WL = White skin lesions

**Supplementary Table 2.** Primers used to identify specific virus sequences in individual bats.

| Name | Sequence (5'-3') | Virus target | Gene target | Fragment size (bp) |
|---|---|---|---|---|
| **BRV_gag_F1** | ACCTTCAATCTCCCTGTCAT | Hervey pteropid | gag | 483 |
| **BRV_gag_R1** | AAAAGGCCAATATTGTAGGG | gammaretrovirus | | |
| **BRV_pol_F2** | ATGTGACTGTGATTGCTTCC | | pol | 444 |
| **BRV_pol_R2** | TATTGATGTCCTTTCCATCG | | | |
| **BRV_env_F3** | GCGGACCAACTATTGTGATA | | env | 421 |
| **BRV_env_R3** | CAGAGATGTTGGTGGGTAGA | | | |
| **Bat_pegi_F1** | TGTACACCACATTTCACTCC | Bat pegivirus | NS3 | 571 |
| **Bat_pegi_R1** | GTGAGCTTCTTCATGTACGG | GHFF04/Li/1 | | |
| **Bat_pegi_F2** | GTTGGTTGAGGCATGTGTTG | | NS5B | 211 |
| **Bat_pegi_R2** | GTAGGCGGGTCCCATAATTT | | | |
| **Kunsagi_F1** | CTTCCGTATGCAGTTCTTGGA | Auskunsag virus | 3D | 397 |
| **Kunsagi_R1** | TGCGTCTGGATTACACTTGAG | | | |

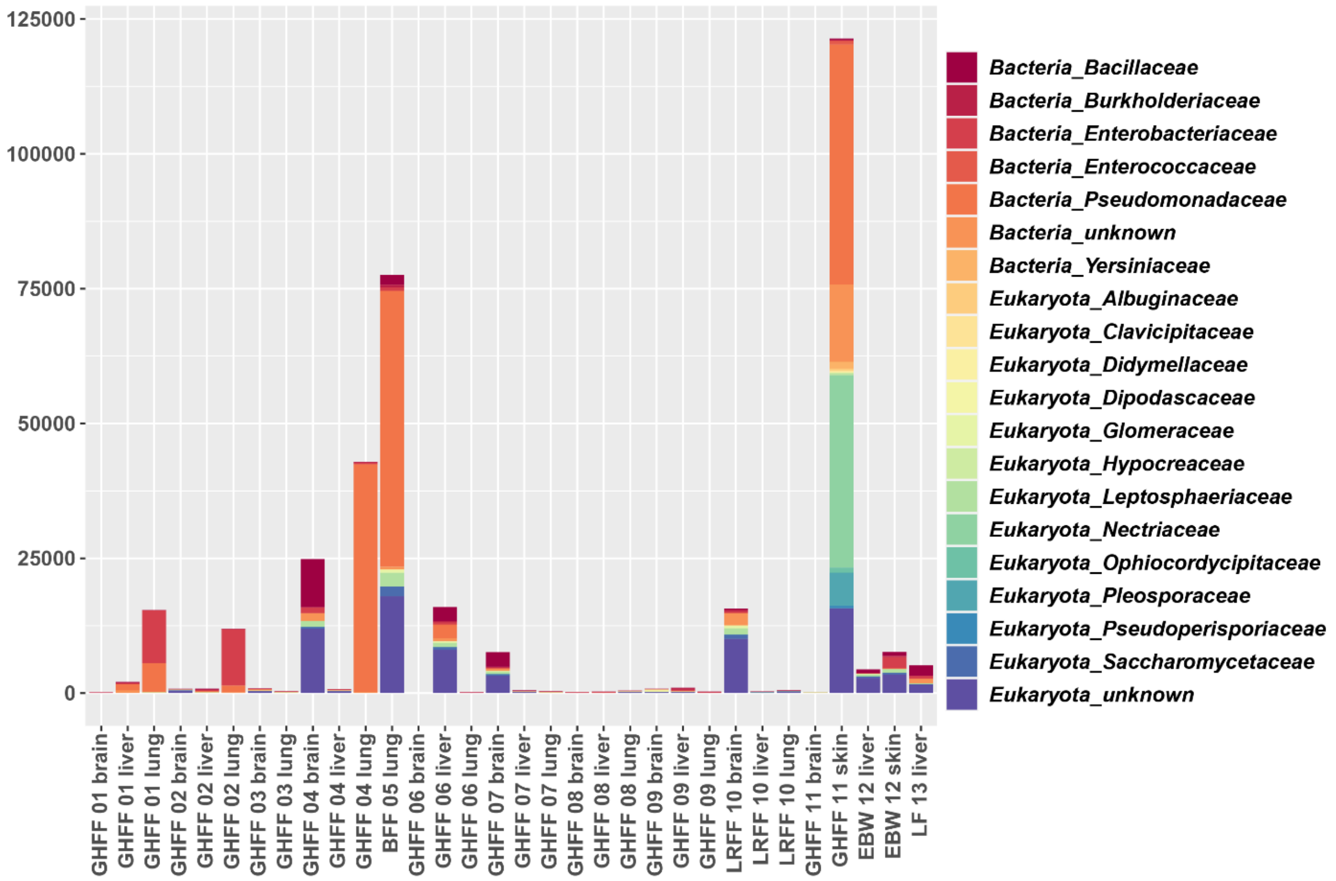


**Supplementary Figure 1** Bacterial and fungal composition within lung, brain, liver and skin samples. Columns represent the read abundance for each family presented as RPM. Only the top 20 families are displayed in the graph.
